# Honey bee queen susceptibility to viral infection varies across developmental stages in queen rearing operations

**DOI:** 10.1101/2025.10.13.682001

**Authors:** Sagar Bhandari, Somayeh Mehrparvar, Omar Martinez Caranton, Bita Valizadeh, Julie Hardy, Michael Simone-Finstrom, Jeffrey W. Harris, Esmaeil Amiri

## Abstract

Viruses are a large class of honey bee pathogens that negatively affect colony health, yet their prevalence and transmission dynamics in commercial queen production operations remain poorly understood. To address this gap, we conducted a series of controlled queen monitoring experiments and surveys to understand the prevalence and viral loads of seven viruses across developmental stages of queens, drones, royal jelly, and workers from associated colonies. All viruses except SBV were detected, with BQCV, DWV-B, and LSVs showing consistently high prevalence. Eggs were frequently infected with LSVs, DWV-B, and CBPV, suggesting vertical virus transmission, and highlighting the importance of selecting healthy breeder queens. BQCV, on the other hand, dominated in queen larvae, pupae, and adult stages. Mated queens, particularly those maintained in bank colonies, exhibited higher prevalence and viral loads than virgin queens, with DWV-B and BQCV being most abundant. Worker bees from bank colonies also showed slightly higher viral loads compared to other colonies, indicating potential risks associated with queen banking. Drone samples revealed high BQCV and DWV-B prevalence, indicative of their potential role in venereal transmission. The results from hierarchical clustering and correlation analyses provided evidence that viral profiles of queens did not necessarily match those of their resident colonies, highlighting complex viral transmission dynamics. Collectively, these findings provide novel insights into virus transmission dynamics during queen production and emphasize the need to improve queen health.

**Author summary:** Honey bee queen failure is a frequent challenge in beekeeping operations, and viral infections are increasingly recognized as an important contributor. Since no chemical therapeutics are available for managing viral diseases in honey bees, understanding routes of viral transmission is critical for developing strategies to minimize infections. However, the diversity of viruses infecting queens, their prevalence, and their main transmission routes have not been systematically investigated within the queen production industry. Therefore, in this study we investigated viral prevalence, loads, and transmission dynamics in commercial queen production operations, using a diverse array of samples, including queens at different developmental stages, worker bees from associated colonies, drones, and *Varroa* mites. Our analysis consistently indicates that vertical transmission from infected breeder queens, natural mating, and bank colonies with high *Varroa* infestations are key points where queens are most vulnerable and become infected with viruses. These findings offer novel insights into virus transmission dynamics in queens, emphasizing the importance of selecting healthy breeder queens and controlling *Varroa* infestations in drone source and bank colonies.

## Introduction

As the sole reproductive female in the colony, a honey bee (*Apis mellifera*) queen lays eggs to replenish old worker bees [1] and releases pheromones to regulate various colony functions [1–4]. Therefore, the health and reproductive vigor of the queen have direct impacts on the performance of the colony she leads [5–7]. Pathogens can negatively impact queen performance by mediating trade-offs among key physiological and reproductive traits [8–10]. Among them, viruses represent a large class of honey bee pathogens that can infect queens at different developmental stages [8,11,12], and adversely impact reproduction capacity, pheromone profile, and consequently their lifespan [8,10,13,14]. Overt viral infections are rarely observed in honey bee queens in colonies, as infected queens, regardless of their life stage, are often removed by worker bees before clinical symptoms become noticeable [15]. On the other hand, covert viral infections typically do not cause observable symptoms in queens [13], yet they may compromise the quality of young queens by impairing their reproductive potential and contributing to queen supersedure [10].

Several honey bee viruses, including deformed wing virus (DWV), black queen cell virus (BQCV), sacbrood virus (SBV), Kashmir bee virus (KBV), and acute bee paralysis virus (ABPV), have been detected in very young queen larvae [12,16–18]. It has also been demonstrated that infection levels in queen larvae were significantly higher compared to those in eggs and pupae [12]. These infections in developing queens can originate either through vertical transmission from parents [17,19,20] and/or horizontally from nurse worker bees through infected food from glandular secretions, or body contact [13,21]. Unlike worker bees and drones, queens are generally not infected by *Varroa* mites, except in the rare case when no worker or drone brood is available in the colony or when the colony is highly infested with mites [22]. The consumption of virus particles in brood food by queen larvae can lead to infection and may result in larval and pupal mortality if a sufficiently large number of virus particles are ingested, as observed in the case of BQCV [23]. In addition, the accumulation of virions (BQCV, DWV, and Israeli acute paralysis virus (IAPV)) in the wax of queen cells may infect the late larval or early pupal stages when a queen’s body contacts the wax [24]. A few studies have sporadically detected different viruses in newly emerged queens [12,18], but mated queens have regularly been found to be infected with several viruses and high virus titers [8,25,26]. Infection in mated queens can occur through mating due to venereal transmission [27,28] or through horizontal transmission pathways, either through body contact when housed with infected worker bees or through trophallaxis if they receive food from infected workers [21,29].

In commercial queen production operations, banking queens is a routine procedure, which is basically the storage of queens by caging them individually and placing them into specially prepared strong colonies, called queen banks [30]. Queen banking offers several benefits to queen producers. It frees up mating colonies to increase production of naturally mated queens during early spring. Queen producers can also bank a surplus of queens throughout the summer to requeen colonies later in the season. Additionally, mated queens can be banked over winter to ensure the availability of queens when making splits or requeening colonies in the early spring [31]. However, potential unfavorable environments inside banks, with virus-infected old worker bees in the bank, combined with the confinement of queens, may act as stressors that make queens susceptible to viral infections [30,32]. Therefore, the impact of banking on virus transmission and infections in queens needs to be evaluated.

Although there are a few direct options for beekeepers to mitigate virus infections in their colonies [33], none of these potential treatments are practical in typical queen production systems. Therefore, identification of critical developmental stages at which queens are most vulnerable to viruses and transmission routes is vital to the development of approaches to minimize or interrupt viral infections in honey bee queens. Transmission routes provide insights into a pathogen’s virulence, spread, persistence, and evolutionary relationships with its host [17,34]. So far, most studies of viral transmission in honey bee queens have been limited to a single transmission pathway, thereby overlooking the complexities of virus transmission [19,27,28,35]. A comparative analysis of the effectiveness of these routes in transmitting viruses, particularly in real-world queen production operations, is essential for developing effective management strategies. To address these concerns, a series of surveys and controlled queen monitoring experiments were conducted to identify the stages during the queen production process when queens are most susceptible to viral infections. Queen-destined individuals were analyzed throughout the progression from eggs in breeder colonies to larval and pupal development in nurse colonies to mated queens in banked colonies, while also considering the roles of workers and drones in the transmission of viruses.

## Results

### Virus Prevalence in Commercial Queen Production Operations

The prevalence of seven honey bee viruses was analyzed across different developmental stages of queens, and worker bees in different colony types. Virus prevalence was calculated as the percentage of samples with a detectable viral load (i.e., viral load > 0). Among the tested viruses, all of them, except SBV, were detected in every developmental stage; however, there was a notable variation in the prevalence of these viruses within and across the stages (Fig 1A). Eggs were mostly infected with LSVs (71.8%), DWV-B (66.9%), and CBPV (51.61%), suggesting vertical transmission. However, in the case of the queen larvae and pupae, BQCV was predominantly detected at 55.67% and 30.69%, respectively. Royal jelly exhibited a lower overall virus prevalence, with BQCV (14.6%) and DWV-A (3.12%) being comparatively the most prevalent. Among the various developmental stages of the honey bee queens, the pupal stage exhibited the lowest virus prevalence, with all detected below 31%. In virgin queens, BQCV showed the highest prevalence (37.08%), while CBPV (31.46%) was also frequently detected. BQCV (87.5%), DWV-B (37.5%), and CBPV (25%) were dominant in mated queens sampled from mating nucs, whereas the rest of the viruses were detected below 10%. Notably, BQCV (95.19%) and DWV-B (75.85%) had the highest prevalence in mated queens in banks, among other stages. The prevalence of IAPV and SBV was consistently low, with less than 5% detection across stages.

**Fig 1.**
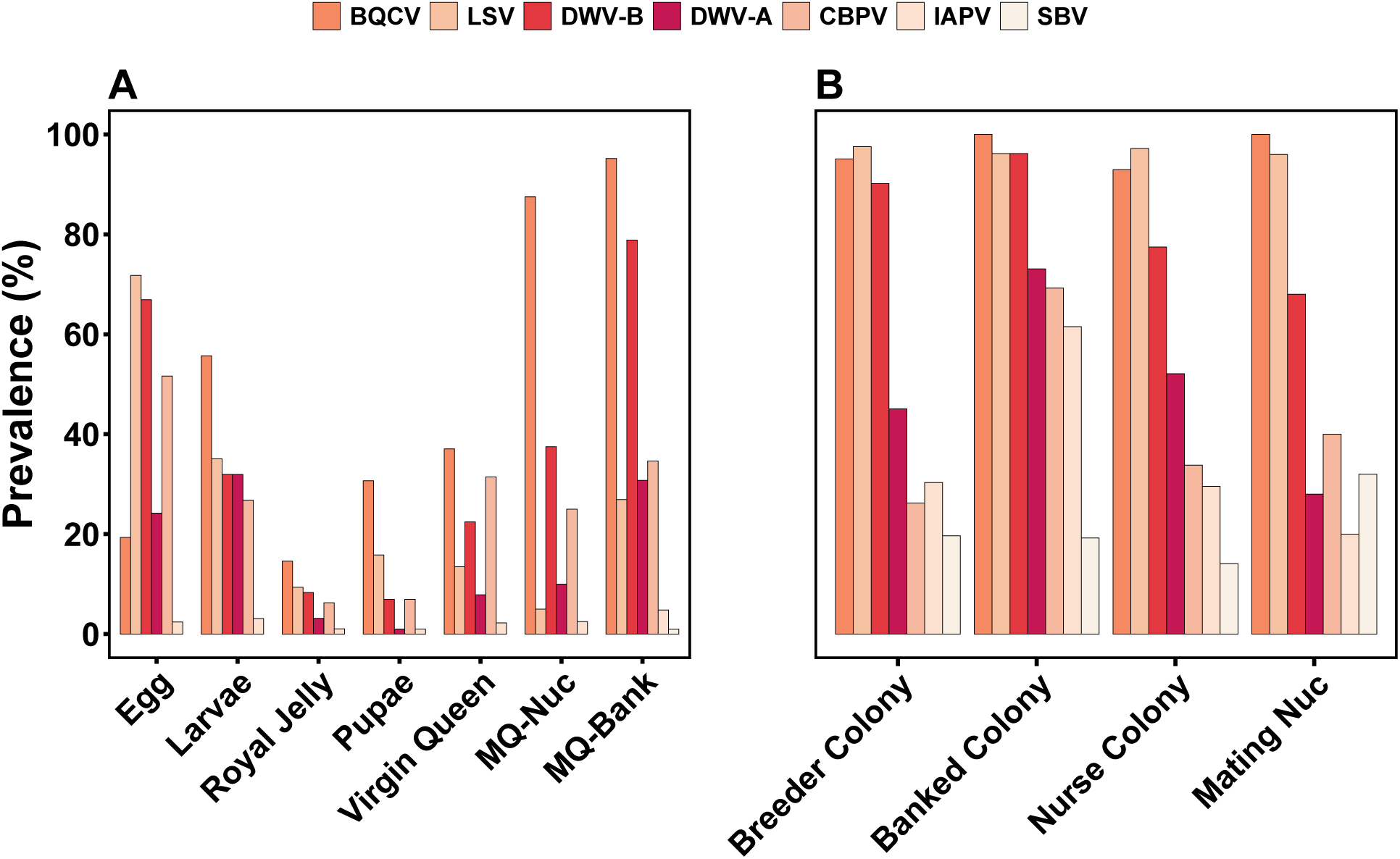
Virus prevalence (%) in honey bee queens across developmental stages and in worker bees from different colony types. (A) Most of the viruses showed high prevalence in the early developmental stages (eggs and larvae). The prevalence is reduced in pupae and virgin queens, and spikes upward in the adult queens, particularly after mating. MQ-Nuc refers to mated queens that were sampled from mating nucs, and MQ-Bank refers to mated queens that were sampled from queen bank colonies. (B) Across the colony types, BQCV, LSV, and DWV-B showed high prevalence (>68%), DWV-A and CBPV showed moderate to high prevalence (26-73%), and IAPV and SBV showed low to moderate prevalence (14-61%) compared to other viruses. The banked colony showed a relatively higher prevalence for most of the tested viruses.

Variations of virus prevalence were also observed in different colony types that are used in queen rearing (Fig 1B). All seven tested viruses were found to be prevalent in each colony type; however, BQCV, DWV-B, and LSVs were among the highly prevalent viruses at the colony level, reported in >68% of the tested colonies. Among these three highly prevalent viruses, BQCV and LSVs had a similar prevalence (>90%) irrespective of the colony types; however, a fluctuation in the prevalence of DWV-B was observed, showing the highest prevalence in banked colonies (96.2%) and the lowest prevalence in mating nucs (68%). Banked colonies exhibited the highest prevalence of five viruses: BQCV (100%), DWV-B (96.2%), DWV-A (73.08%), CBPV (69.23%), and IAPV (61.54%), suggesting potential risks associated with queen banking. Except for banked colonies, the detection of CBPV, IAPV, and SBV was low (<40%) in other colony types. LSVs showed the highest prevalence in breeder colonies (97.54%), and SBV in mating nucs (32%). Though most of the viruses were highly prevalent in nurse bees, their prevalence in royal jelly was very low or absent. Among the DWV variants, DWV-B was highly prevalent in both the developmental stages and colony types than DWV-A.

Additionally, drones were sampled individually from each operation while returning from their mating flights for both years. BQCV showed the highest prevalence (85.47%), followed by DWV-B (70.39%), and LSVs (42.46%). The prevalence of DWV-A (31.28%) and CBPV (30.73%) was similar; however, less than 10% of drones were positive for IAPV and SBV (S4 Table).

### Viral loads in commercial queen production operations

The viral load data from both years were combined and are presented in two subsets: one representing the viral load in different developmental stages of queens and another representing the viral load in worker bees from different colony types. DWV-A, DWV-B, BQCV, LSVs, and CBPV were mostly prevalent in the samples; therefore, the viral loads are presented for these viruses (Fig 2).

**Fig 2.**
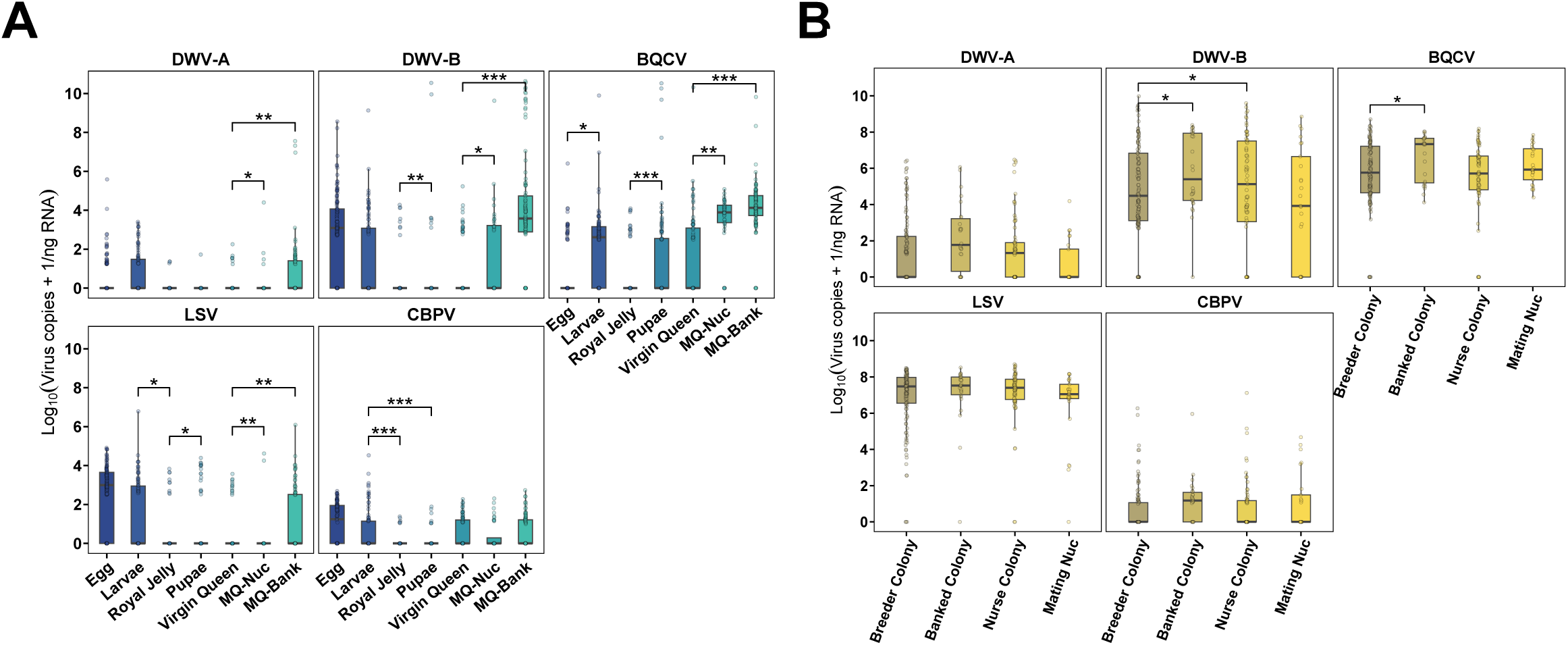
Viral loads in honey bee queens at different developmental stages and in worker bees from different colony types in the survey. (A) Larvae, royal jelly, pupae, and virgin queens were from the nurse colonies. MQ-nucs refers to mated queens in mating nucs, and MQ-bank refers to mated queens in bank colonies. Across developmental stages, viral loads were similar in the early stages for most viruses; however, they increased after mating, as mated queens consistently exhibited higher viral loads, except for CBPV, than virgin queens. (B) The viral load across colony types varied only for DWV-B and BQCV, as banked colonies exhibited higher viral loads than breeder colonies for both viruses. In all panels, the black horizontal lines in boxplots represent the median of the (log_10_ (viral copies + 1)) transformed viral load, and each dot represents the viral load in the particular sample type. Significance levels based on Bayesian posterior distributions are indicated as: p < 0.05 (*), p < 0.01 (**), and p < 0.001 (***) as estimated via Bayesian pairwise comparisons.

#### Viral loads in different developmental stages of queens

To investigate the viral loads across the developmental stages of queens, a pairwise comparison was made between eggs and larvae, larvae and pupae, larvae and royal jelly, royal jelly and pupae, virgin queens and mated queens sampled from mating nucs, virgin queens and mated queens sampled from bank colonies, and mated queens from mating nucs and the ones sampled from bank colonies (Fig 2A). Variation in viral loads was observed across different developmental stages (eggs, larvae, pupae, virgin queens, and mated queens) of queens and royal jelly; however, the pattern of viral loads was similar for multiple viruses. Eggs and queen larvae exhibited similar viral loads for all the viruses except for BQCV, in which eggs had significantly lower BQCV loads than larvae (median ratio=0.225, 95% CI: 0.056–0.91, p=0.022). Viral loads in larvae and pupae differed only for CBPV, as larvae exhibited higher CBPV load compared to pupae (median ratio=24.222, 95% CI: 6.656–92.605, p<0.001). However, royal jelly had significantly lower viral loads than larvae and pupae for most of the viruses. For instance, significantly higher viral loads were observed in larvae than in royal jelly for LSVs (median ratio=14.35, 95% CI: 4.04–54.14, p=0.026) and CBPV (median ratio=41.285, 95% CI: 9.572-187.783, p<0.001). Similarly, royal jelly had significantly lower viral loads than pupae for DWV-B (median ratio=0.02, 95% CI: 0.0003–1.39, p<0.01), BQCV (median ratio=0.063, 95% CI: 0.01–0.39, p<0.001), and LSVs (median ratio=0.25, 95% CI: 0.06–0.99, p=0.032). Pupae had comparatively higher viral titers for DWV-B and BQCV, with values > 10 (log_10_ (virus copies+1)/ng RNA) in some samples. After mating, the viral load increased significantly in queens for all viruses except CBPV. Virgin queens had significantly lower viral loads than mated queens from mating nucs for DWV-A (median ratio=0.086, 95% CI: 0.0021–3.36, p<0.05), DWV-B (median ratio=0.13, 95% CI: 0.007–2.367, p<0.05), BQCV (median ratio=0.165, 95% CI: 0.037–0.72, p=0.005), and LSVs (median ratio=0.096, 95% CI: 0.007–1.376, p=0.008). Similarly, the viral load was lower in virgin queens compared to mated queens from bank colonies for DWV-A (median ratio=0.025, 95% CI: 0.002– 0.334, p<0.01), DWV-B (median ratio=0.009, 95% CI: 0.001–0.0772, p<0.001), BQCV (median ratio=0.093, 95% CI: 0.027–0.313, p<0.001), and LSVs (median ratio=0.216, 95% CI: 0.064– 0.705, p=0.007). Across all the viruses, there was no significant difference in viral loads between mated queens from mating nucs and mated queens from bank colonies. However, viral loads differed more strongly between virgin queens and mated queens from bank colonies than between virgin queens and mated queens from mating nucs for DWV-A, DWV-B, and BQCV, as indicated by the p-values. In general, the DWV-B titer was higher in all the developmental stages of queens than DWV-A, but the infection patterns of DWV-B were similar to DWV-A. Across all developmental stages, DWV-A and CBPV titers were consistently low, with values below 2 (log_10_ (virus copies + 1)/ng RNA).

#### Viral loads in worker bees from different colony types and drones

To understand the virus infection and transmission at the colony level, worker bees were collected from breeder colonies, nurse colonies, bank colonies, and mating nucs, and pairwise comparisons of viral loads were made among them (Fig 2B). Significant differences in viral loads among colony types were observed only for DWV-B and BQCV. Worker bees in breeder colonies exhibited significantly lower DWV-B loads than banked colonies (median ratio=0.145, 95% CI: 0.0225– 0.948, p<0.05) and nurse colonies (median ratio=0.219, 95% CI: 0.0564–0.828, p<0.05). Similarly, worker bees in breeder colonies showed significantly lower BQCV loads than those in the banked colonies (median ratio=0.328, 95% CI: 0.099–1.059, p=0.05). DWV-A and CBPV exhibited consistently low viral titers across colony types with a median of <2 (log_10_ (virus copies+1)/ng RNA). In contrast, LSVs exhibited consistently high viral titers across colony types, with a median of around 7 (log_10_ (LSV copies + 1)/ng RNA), showing no significant differences. Additionally, the mean log_10_ (virus copies + 1)/ng RNA in drones was 0.72, 3.37, 4.0, 1.67, and 0.55 for DWV-A, DWV-B, BQCV, LSVs, and CBPV, respectively. Detailed statistics on drones are presented in the S4 Table.

### Viral profiling in queens and associated colonies

Hierarchical cluster analysis was performed to identify patterns of viral loads across queen developmental stages and colonies in the survey, revealing three clusters (Fig 3). The young developmental stages (larvae, pupae, and virgin queens) formed one cluster. However, mated queens from both bank colonies and nucs, along with eggs, exhibited different viral load profiles than other stages and clustered separately. The distinct clustering of eggs from subsequent developmental stages (larvae, pupae, and virgin queens) suggests the existence of additional transmission routes alongside vertical transmission. Similarly, worker bees from nurse colonies, breeder colonies, bank colonies, and nucs clustered together along with royal jelly. Among them, two subclusters: one between breeder colonies and bank colonies, and the other between mating nucs and royal jelly, and a branch representing nurse colonies, can be observed. Though nurse bees and royal jelly exhibited somewhat similar infection patterns, the viral load in royal jelly was very low. Moreover, different infection patterns in mated queens and worker bees from mating nucs suggest unrelated viral load profiles. Regarding viruses, BQCV and the rest of the viruses broadly formed two clusters. Among the other viruses, CBPV and LSVs clustered together, which are first closely associated with DWV-A and then with DWV-B.

**Fig 3.**
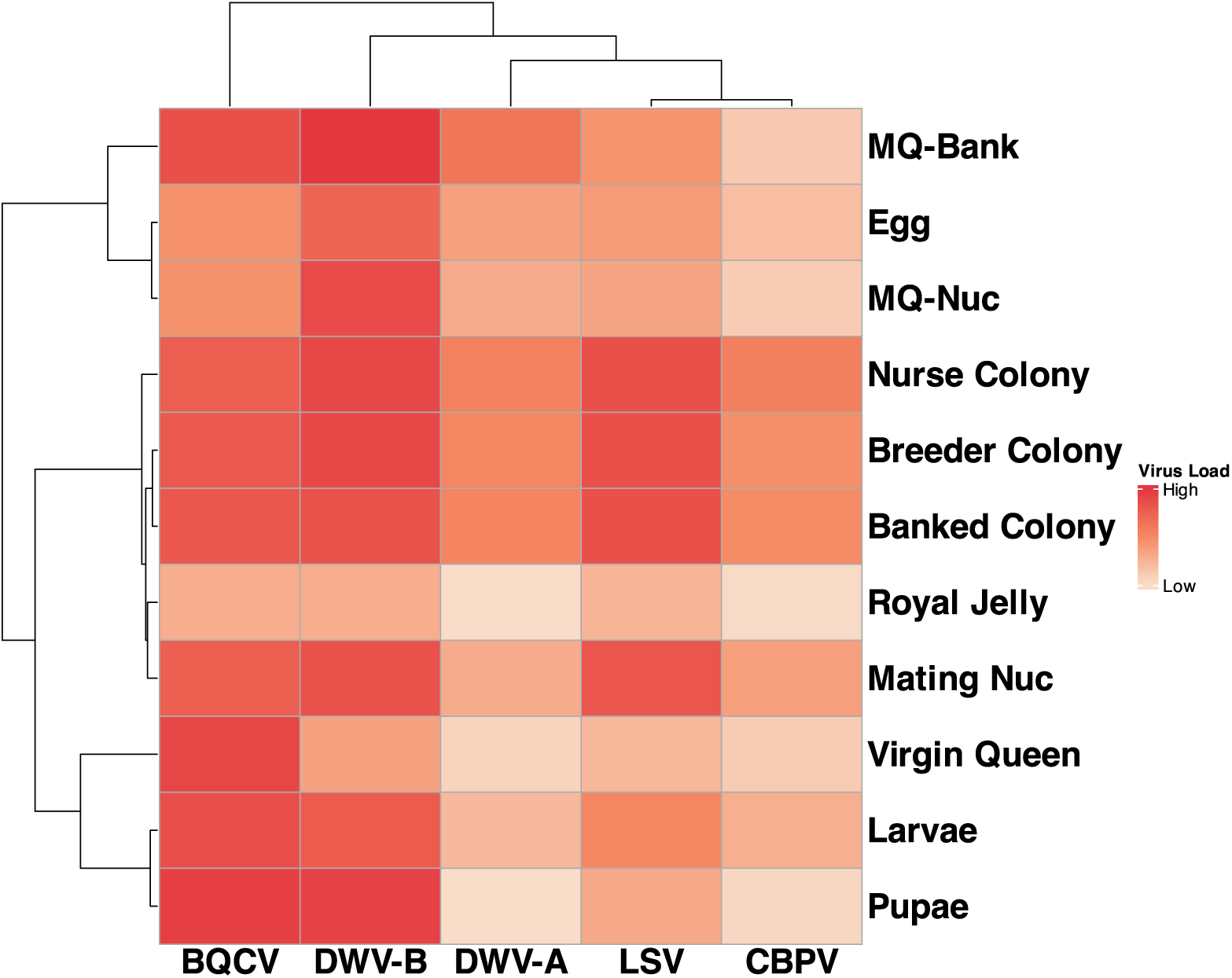
Hierarchical cluster analysis of viral loads in developmental stages and colony types. MQ-nucs refers to mated queens in mating nucs, and MQ-bank refers to mated queens in bank colonies. BQCV and DWV-B showed high viral loads, DWV-A and LSV showed moderate to high viral loads, and CBPV showed low viral loads across developmental stages. However, BQCV, DWV-B, and LSV showed high viral loads, and DWV-A and CBPV showed moderate to high viral loads at the colony level. The viral loads across all the samples follow the order: DWV-B > BQCV > LSV > DWV-A > CBPV.

### Correlation analysis

We specifically explored correlations between viral loads in royal jelly and those in the corresponding queen larvae, as well as between mated queens from mating nucs and the corresponding worker bees collected during the survey (Table 1). Since these were the only corresponding sample pairs with known viral loads, correlation analysis for viral loads was limited to these comparisons. Royal jelly and queen larvae showed a positive correlation for BQCV, DWV-A, DWV-B, and LSVs; however, the correlation was significant only for DWV-A (ρ=0.296, p=0.003) and DWV-B (ρ=0.377, p=1.6e-04). Similarly, a weak correlation was found between mated queens and the corresponding worker bees from mating nucs without any significant difference, indicating unrelated viral load profiles between them.

**Table 1.** Correlation analysis between viral loads in royal jelly and larvae, and between mated queens and worker bees from mating nucs.

| Viruses | Royal jelly and larvae |  | Queens and worker bees from mating nucs |  |
| --- | --- | --- | --- | --- |
|  | Correlation coefficient | p-value | Correlation coefficient | p-value |
| BQCV | 0.176 | 0.086 | -0.218 | 0.294 |
| CBPV | -0.155 | 0.132 | 0.198 | 0.343 |
| DWV-A | 0.296 | 0.003 | 0.079 | 0.706 |
| DWV-B | 0.377 | 1.6e-04 | -0.248 | 0.231 |
| LSVs | 0.019 | 0.851 | 0.062 | 0.768 |

While significant differences were found in mite infestation levels across different colony types (significantly lower in breeder colonies than nurse colonies, p=0.0127) (S1 Fig), analysis of paired samples indicated no significant relationship between viral load and mite levels (S2 Fig and S3 Fig). This was likely due to a lack of high range in *Varroa* levels across the samples that could be compared.

### Viral loads in the controlled experiment

In this experiment, viral infections were actively monitored throughout the queen production process, and viral loads are presented in two subsets. The first one represents viral loads in the mother queen and different developmental stages of daughter queens (Fig 4A). The second subset represents viral loads in eggs and worker bees from mating nucs sampled at two different time points, and worker bees from nurse colonies (Fig 4B). We did not detect IAPV and SBV in most of the samples; therefore, viral loads are presented for DWV-A, DWV-B, BQCV, LSVs, and CBPV.

**Fig 4.**
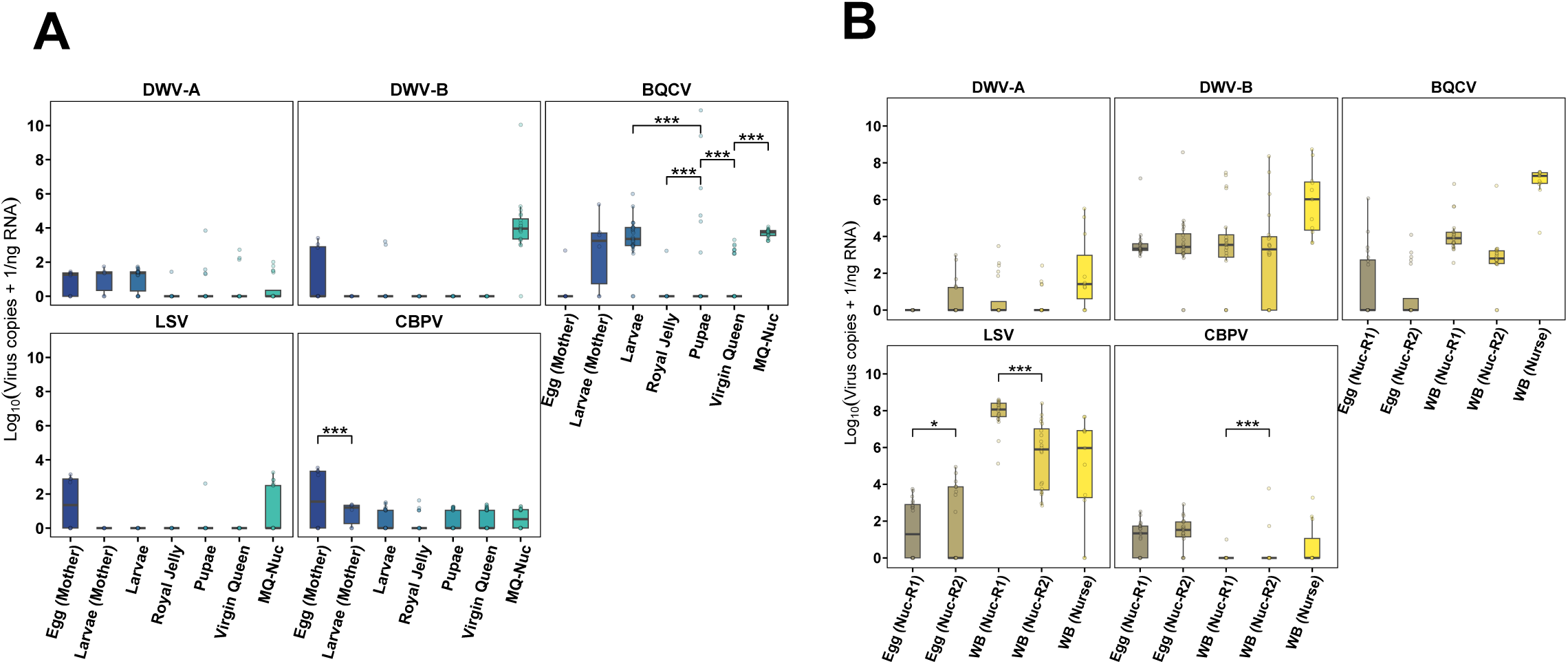
Viral loads in honey bee queens at different developmental stages and in worker bees from different colony types in the controlled experiment. (A) Egg (mother) and larvae (mother) refer to the eggs and larvae in the mother or breeder colony, respectively. Larvae, royal jelly, pupae, and virgin queens were from the nurse colonies. MQ-nucs refers to mated queens in mating nucs. (B) Egg (Nuc-R1) and Egg (Nuc-R2) refer to eggs collected from mating nucs at the first and second rounds at an interval of 6 weeks. Similarly, WB (Nuc-R1) and WB (Nuc-R1) refer to worker bees collected from mating nucs at the first and second rounds at an interval of 6 weeks. WB (Nurse) refers to worker/nurse bees collected from nurse colonies. In all panels, the black horizontal lines in boxplots represent the median of the (log_10_ virus copies + 1) transformed viral load, and each dot represents the viral load in the sample type. Significance levels based on Bayesian posterior distributions are indicated as: p < 0.05 (*) and p < 0.001 (***) as estimated via Bayesian pairwise comparisons.

#### Viral loads in the mother queen

Eggs and very young larvae were collected from the mother colony to assess the virus infection status of the mother queen. All tested viruses were detected from egg samples, with a median of virus titers below 2 (log_10_ (virus copies+1)/ng RNA) for all the viruses (Fig 4A). However, larvae were infected only with DWV-A, BQCV, and CBPV, in which BQCV exhibited comparatively higher viral loads with median log_10_ (BQCV+1)/ng RNA>3. No significant difference was observed between eggs and larvae for all viruses except for CBPV, wherein eggs had higher CBPV titers compared to larvae (median ratio=74.176, 95% CI: 18.171-307.007, p<0.001).

#### Viral loads in different developmental stages of queens

Across queen developmental stages (larvae, pupae, virgin queens, and mated queens), the infection pattern was similar, with no statistical differences for all the viruses except for BQCV (Fig 4A). While queen larvae were infected with all viruses tested except LSVs, BQCV exhibited higher viral titers with median log_10_ (BQCV+1)/ng RNA>3. Royal jelly was positive for DWV-A, BQCV, and CBPV, but the virus titers were low (median<2 log_10_ (virus copies + 1)/ng RNA). Although both larvae and pupae showed elevated BQCV loads, larvae exhibited BQCV loads lower than pupae (median ratio=0.014, 95% CI: 0.001-0.125, p<0.001). Similarly, pupae showed higher BQCV loads than virgin queens (median ratio=3.29 × 10³, 95% CI: 238.151-4.52 × 104, p<0.001). Additionally, lower BQCV titers were observed in virgin queens compared to mated queens from mating nucs (median ratio=0.018, 95% CI: 0.003-0.135, p<0.001). As DWV-B and LSVs were not detected in virgin queens, the comparison between virgin and mated queens is statistically inconclusive for these viruses. Across all developmental stages, DWV-A, LSVs, and CBPV consistently exhibited low viral loads (median<2 log_10_ (virus copies + 1)/ng RNA).

#### Viral loads in queens after mating

To assess the viral loads in queens after mating, eggs were collected from mating nucs twice at an interval of six weeks along with corresponding worker bees. As reported in the mother colony, eggs were infected with all the viruses; however, the viral titers in eggs differed only for LSVs when collected six weeks apart (Fig 4B). The first round of eggs had significantly lower LSVs titers than the second round (median ratio=0.155, 95% CI: 0.013-2.058, p=0.037). Additionally, worker bees showed differences in virus titers for LSVs and CBPV when collected at the same interval. The LSVs copy number in the first round of worker bees was higher than that in the second round (median ratio=213.531, 95% CI = 41.233 - 1.13 × 10³, p<0.001). In contrast, worker bees collected in the first round showed substantially lower CBPV loads as compared to the second round (median ratio=0.009, 95% CI: 0-0.159, p<0.001). Additionally, worker bees were collected from nurse colonies to assess colony-level infection and virus transmission to royal jelly. Although nurse bees were highly infected with DWV-B, BQCV, and LSVs, with a median of more than 5 log_10_ (virus copies + 1)/ng RNA, the virus titer in royal jelly was very low or absent.

## Discussion

Our study clearly demonstrates variation in the prevalence and titers of viruses across the developmental stages of queens and associated colonies in queen production operations. The survey results indicate that all tested viruses (except SBV) were detected in eggs collected from breeder colonies, suggesting vertical transmission of these viruses [12,17,20,36,37]. However, there were notable variations in the incidence and titers of these viruses in eggs. As reported in our case, the high incidence and titers of DWV-B, BQCV, and LSVs in eggs have previously been documented [12,37]. Our findings of a high prevalence and low titer of CBPV are consistent with [38], who also reported low CBPV genomic load in eggs, larvae, and pupae below 2×10^3^ CBPV copies per individual. These findings reveal a pattern of a low-level, widely persistent, and less virulent nature of this virus, particularly in the early developmental stages [39,40]. Although the median viral load was low, a significantly higher CBPV load in eggs compared to larvae was reported in the controlled experiment, suggesting that vertical transmission of CBPV either doesn’t lead to infection or those larvae that do get infected are removed by workers [41].

Additionally, pathogens often reduce their own fitness and appear less virulent without limiting the host’s reproductive capacity during vertical transmission [42,43]. When vertical transmission occurs through biparental routes, as seen in honey bees [19], the transmission efficiency might be further enhanced [44], which explains the high incidence of most viruses in eggs. Vertical virus transmission via eggs may affect later developmental stages when conditions are suitable [12], making it a critical stage in queen production. Since virus levels in eggs can accurately reflect the infection status of queens [12], screening eggs for virus levels is vital for selecting healthy breeder queens and breaking the cycle of vertical transmission.

Cluster analysis showed similar infection patterns in queen larvae and pupae, with a high prevalence of BQCV (55.67% in larvae and 30.69% in pupae) and high viral loads of BQCV and DWV-B. High prevalence and titers of BQCV and DWV-B in queen larvae and pupae were also reported in previous studies [12,35,45], which indicates that these stages are suitable for these viruses to replicate [46]. It has been reported that BQCV was the most common reason behind queen larvae mortality [47], and thus, it is a virus of economic importance for queen producers. In the survey, queen larvae exhibited significantly higher BQCV loads than eggs and higher CBPV loads than pupae. Domingues et al. [12] reported that viral loads of BQCV, DWV-A, LSVs, and SBV in larvae were significantly higher than in eggs and pupae. Our study could not answer whether the virus replication in larvae was higher during the early instars or the later instars. Therefore, further investigation is required to assess the temporal variation of viral load in queen larvae to identify the critical infection window.

Among the developmental stages, queen pupae exhibited the lowest prevalence of the five viruses (BQCV, DWV-B, DWV-A, CBPV, and IAPV), with all viruses present at a prevalence below 31%. In contrast, higher BQCV loads in pupae than larvae in the controlled experiment suggest that the pupal stage is equally or even more susceptible to BQCV infection than other stages. The detection of BQCV in dead queen prepupae and pupae supports this finding [23]. Nevertheless, the overall viral incidence was lower in pupae, which might be due to several possible reasons. One possible explanation is a lack of feeding during the pupal stage. For instance, studies conducted in another social insect, *Solenopsis invicta*, demonstrated higher loads of IAPV and Solenopsis invicta virus 2 in larvae compared to pupae, which was attributed to frequent trophallaxis activity between larvae and worker ants [48,49]. Another plausible reason might be the physiological changes at larval-pupal metamorphosis. Results from another holometabolous insect, *Bombyx mori*, reported that as pupal duration increased, pupae exhibited greater resistance to nuclear polyhedrosis virus infection [50]. During larval-pupal metamorphosis in *Apis mellifera*, the larval midgut epithelium degenerates, and regenerative cells actively proliferate, forming a new midgut layer in pupae [51]. The histolysis of larval midgut epithelium and the gradual reduction in the number of cellular receptors for viruses during pupal development may be the reason for reduced viral susceptibility during this stage [50]. Furthermore, we cannot exclude the possibility that virus-infected larvae or pupae may die or fail to complete the pupal stage, or be removed by workers, resulting in the inclusion of only healthy or lowly infected pupae in our samples. However, the death of the larval or pupal stage also depends on the virus with which they are infected. For instance, high pupal mortality was reported before adult eclosion when insects were injected with high levels of BQCV and SBV [52]; on the other hand, mortality was low or absent when insects were injected with DWV [52,53]. Therefore, further investigation is required to determine the exact reason behind the low virus prevalence in queen pupae.

As reported in the cluster analysis, distinct infection patterns in eggs and subsequent developmental stages (larvae, pupae, virgin queens) suggest that, in addition to vertical transmission, some degree of horizontal transmission might play a role. Comparatively higher prevalence and titers of BQCV, DWV-B, and LSVs were observed in young developmental stages (larvae and pupae) and royal jelly, in addition to a positive correlation between larvae and royal jelly, suggesting that these viruses can be transmitted horizontally through trophallaxis activity. A comparable study by Žvokelj et al. [18] reported that when nurse bees were positive for DWV and ABPV, neither of these viruses was detected in royal jelly or larvae. However, when nurse bees were positive for BQCV, the same virus was detected in royal jelly and larvae. On the other hand, royal jelly exhibited lower BQCV loads than pupae in the controlled experiment, while in the survey, a similar relation was observed for DWV-B, BQCV, and LSVs loads. Similarly, the LSVs load was lower in royal jelly compared to larvae. While nurse bees exhibited higher loads for DWV-B, BQCV, and LSVs in both the controlled experiment and the survey, the viral loads in the corresponding royal jelly were very low. The low prevalence and titers of viruses in royal jelly, as well as the weak correlation between viruses in larvae and royal jelly, suggest limited virus transmission through feeding. Our findings align with another study, which reported that infected bees can transmit CBPV to their healthy companions through trophallaxis; however, the virus particles were not sufficient to cause disease readily [54]. Other studies have been inconsistent in detecting virus prevalence in royal jelly, with some confirming its presence [16] and others failing to do so [55]. Overall evidence suggests that we cannot completely exclude the possibility of transmission through trophallaxis; however, the primary pathway for infection in the young developmental stages (larvae and pupae) appears to be from the mother queens or vertical transmission. Other studies have also suggested vertical transmission for not detecting viruses in royal jelly, but rather in larvae, which is not typically associated with *Varroa* mite parasitism [11,55].

In adult queens, the sharp increase in prevalence and titers of most viruses—especially for DWV-B and BQCV—in mated queens from both mating nucs and banks compared to virgin queens, indicated that mating is another critical time point for virus infection. All the viruses were detected below 38% in virgin queens, whereas the incidence reached up to 87.5% in mated queens from nucs, and 95.19% from bank colonies for BQCV. In the controlled experiment, virgin queens showed lower BQCV loads than mated queens from nucs. Similarly, virgin queens exhibited lower DWV-A, DWV-B, BQCV, and LSVs loads than mated queens in the survey from both nucs and bank colonies. A higher prevalence of DWV and *Varroa destructor* virus-1 in mated queens than in virgin queens was also observed in a previous study [8]. As drones showed a comparatively higher prevalence and titers of BQCV, DWV-B, and LSVs, mated queens may have acquired viruses from drones during mating. The detection of viruses in the reproductive tissues of drones, such as semen [56], endophallus [28], testis epithelia [57], seminal vesicle, and mucus gland [57,58], and their ability to take a mating flight under virus infection [27] provides evidence of venereal transmission during natural mating [28]. In contrast, similar viral loads of CBPV were observed between virgin and mated queens in both experiments, suggesting its poor venereal transmission.

However, the explanation for higher viral loads in mated queens extends beyond venereal transmission of viruses. The trade-off between reproduction and immune response might be another possible reason. As both mating and reproduction, as well as immunity, are energy-demanding, a trade-off exists between them in several insect species [59–63], including honey bees [64–66]. For instance, a significant negative correlation between sperm viability and lysozyme expression was reported in honey bee queens [65]. Similarly, in the leaf-cutting ant *Atta colombica*, it has been demonstrated that sperm storage and frequency of mating were negatively associated with the encapsulation response 9 days after colony confounding [67]. In contrast, the increased expression of certain immune-related genes was found in mated individuals compared to unmated ones in bumblebees [68]. It may be due to the differences in the mating system between these insect species, as polyandry increases post-copulatory sexual conflict compared to monandry, which ensures paternity at a higher cost to females [69,70]. As there exists a polyandrous mating system in honey bees, queens might exhibit a suppression of immune response after mating. Although the underlying biochemical mechanism behind such a tradeoff is unclear in honey bees, studies conducted in *Tenebrio molitor* and *Drosophila melanogaster* reported that the reduction of immune response after mating is due to induced synthesis of juvenile hormone (JH) from corpora allata [71,72]. Therefore, increased viral loads in mated queens may be attributed to the venereal transmission of viruses and the reproductive immunity tradeoff in queens after mating.

When comparing mated queens from nucs and bank colonies, the prevalence of all tested viruses was higher in mated queens from bank colonies than in those from nucs, suggesting queen banking as another critical time point in queen production. Regarding viral loads, patterns can be noticed showing higher viral loads in mated queens from bank colonies; however, no significant difference was found between these two queen groups. These findings suggest that increasing the banking period may increase the risk of virus infection in banked queens. While the underlying reason behind higher virus prevalence in banked queens is unclear, multiple factors may be involved. Unlike in mating nucs, queens in a bank colony are confined and stored in bulk, restricting their movement and potentially causing stress, which is defined as a perturbation of the homeostatic response to environmental variations [73]. The stress of crowding or confinement was found to decrease the resistance of lepidopteran larvae against invading pathogens [74]. Confined-specific stressors in animals include restricted movement, absence of retreat space, restricted feeding opportunity, and abnormal social groups, as reviewed in [75]. In this regard, banked queens may have a limited interaction with workers for receiving food and grooming, and such social restriction can downregulate the immune and stress response genes [76]. Moreover, since queens are kept in close proximity, they might compete with each other using pheromones to assert dominance, which could be analogous to the pheromonal conflicts seen during queen duels [77]. The cumulative impacts of all these stressors might induce immunosuppression, thereby increasing vulnerability to opportunistic pathogens, including viruses [73].

Worker bees from different colony types were collected and tested to understand their virus infection patterns and potential impacts on the developmental stages of queens. In the controlled experiment, worker bees were collected from nurse colonies and mating nucs, which were positive for DWV-A, DWV-B, BQCV, LSVs, and CBPV over the six-week interval. In addition, a similar viral load (except for LSVs and CBPV) was observed in workers over the period. A previous study also reported the incidence of BQCV and ABPV in attendant workers collected at 2 weeks and 6 weeks after the establishment of the nuclei [18]. Such persistent infection in workers might be due to the constant circulation of the virus through trophallaxis, and/or the stability in the host-virus interaction without fitness cost on the host [49,78]. The establishment of persistent infection in insects is described as the combined action of RNA interference and the reverse transcription of viral RNA, which suppresses the virus replication without complete clearance [79]. In contrast, the variation of LSVs loads in both workers and eggs collected at six-week intervals suggests a seasonal nature of this virus [80–82].

In the survey, all colony types exhibited the prevalence of seven tested viruses, with workers from the banked colonies showing the highest prevalence of BQCV, DWV-B, DWV-A, CBPV, and IAPV. Regarding viral loads, bank colonies showed higher DWV-B and BQCV loads than breeder colonies. In bank colonies, if any queens are infected, they might serve as a reservoir of infection, increasing the possibility of horizontal transmission of the virus through trophallaxis. Due to the presence of multiple queens, there may be less social harmony among workers, which can create stressful conditions, thereby increasing the likelihood of infections. On the other hand, breeder colonies are often selected for hygienic behaviors, and such hygienic workers were reported to exhibit lower *Varroa* mite infestation and DWV loads [83,84]. In our case, we noticed the lowest mite infestation in breeder colonies. The *Varroa* mite has been reported to transmit multiple viruses, including BQCV, DWV-A, DWV-B, LSVs, KBV, and SBV [16,85–87], which may be the reason for lower viral loads in breeder colonies than in bank colonies.

Cluster analysis revealed different patterns of infection between mated queens from mating nucs and the corresponding worker bees. Similar findings were observed in the correlation analysis, with no significant positive correlation between mated queens and the corresponding worker bees in mating nucs. This finding is consistent with Žvokelj et al. [18], who reported that when attendant workers showed infection with ABPV and BQCV six weeks after establishing mating nuclei, the queens in the same nuclei did not show infection with these viruses. When queens were found to be infected with DWV, the attendant workers in the same mating nucleus were free of DWV. It is further supported by different viral load profiles between eggs and breeder colonies reported in our case. These findings suggest that virus profiles in queens do not necessarily match those of their resident colonies [88]. Overall, our results suggest the idea that queens are buffered from viral infection in their colonies, which in turn influences what is transmitted vertically. Since they can’t be buffered from transmission during development or during mating, they don’t match the colonies super well. It is also worth noting that both genotypes of DWV (A and B) exhibited somewhat similar infection patterns across sample types in hierarchical clustering; however, our results are consistent with global trends, where DWV-B is more prevalent and exhibits higher loads than DWV-A [89–92].

In summary, we provided a comparative analysis of various viral transmission routes by testing a diverse array of samples in queen production, such as queen developmental stages, worker bees from all colony types, and drones, while also assessing *Varroa* mite infestation rate. This study demonstrates that viral infections are widespread in commercial queen production systems, with evidence supporting vertical, horizontal, and sexual transmission. The results highlight that honey bee queens are exposed to multiple viruses at different points in their development, with mating and banking being the critical stages for infection. Viral infection during mating is concerning, as infections in drones may compromise semen quality (however, see [66]) and directly introduce viruses into queens during copulation, which could explain the significant increase in viral loads observed after mating. Among all tested viruses, DWV-B and BQCV were consistently the most prevalent and abundant viruses, which need special attention in queen health monitoring programs. Based on our findings, selecting healthy breeder queens, and maintaining drone source colonies and bank colonies with low mites are the suggested interventions to reduce viral transmission in queen production to better safeguard queen health.

## Materials and methods

### Experimental design and sample collection

#### Experiment 1: Survey of commercial queen-rearing operations

To investigate the pattern of virus transmission and determine the susceptible life stages of queens to viral infections, a comprehensive survey was conducted in commercial queen production operations over a two-year period. In this experiment, various samples were collected from commercial queen producers in Mississippi and Louisiana. These queen producers either reside in Mississippi and Louisiana or move their bees from northern states to Mississippi and Louisiana during queen production season, representing a broad range of regions across the U.S. Samples were collected from 8 queen producers in early spring (late March to mid-April) 2023, and from 10 producers during the same period in 2024. In addition to the early spring sampling, we collected samples one more time from four of the Mississippi queen producers in late spring (late May to early June) of both years. In each operation, samples were collected from breeder colonies, nurse colonies, mating nucs, and bank colonies (for terminology, see S1 Table). Samples of eggs (a pool of 25 eggs/queen) and worker bees (∼150) were collected from 5 breeder colonies per operation. The egg samples represent the viral loads of breeder queens, while the worker bees represent the viral load and viral diversity of the colony that they are leading. From nurse colonies (also referred to as cell builders in the industry), 5 queen larvae and their corresponding royal jelly samples were sampled individually. In addition, 5 queen pupae were sampled from the capped queen cells to assess the diversity and load of viral infections across queen developmental stages. To assess the *Varroa* mite infestation and viral load of nurse colonies, pooled samples of adult worker bees were collected from each of the nurse colonies in each operation.

To evaluate viral loads before and after natural mating, 5 virgin queens and 10 mated queens were collected from each operation. Mated queens were collected from both mating nucs and queen bank colonies, 5 each, with corresponding pools of worker bees. Mated queens from mating nucs were collected approximately 2 weeks after mating, as this duration is sufficient to detect virus in queens if the virus is transmitted from drones during natural mating [28]. Mated queens from banks were also collected approximately two weeks after being stored in banks. Additionally, 10 drones were collected in front of drone source colonies while returning from their mating flight to compare the virus profiles with mated queens. Similar sampling procedures were applied in each queen production operation; however, sample size varied slightly among operations, depending on the availability of certain sample types. Altogether, 1074 samples were collected over 2 years for this survey, which comprised 12 sample categories: eggs, queen larvae, royal jelly, queen pupae, virgin queens, mated queens from queen bank colonies and mating nucs, drones, pools of worker bees from breeder colonies, nurse colonies, banked colonies, and mating nucs (S1 Table). All the samples were transported in a cold chain to the laboratory at the Delta Research and Extension Center (DREC), Stoneville, MS, and stored at -80°C until RNA extraction.

#### Experiment 2: Controlled experiment

This experiment was designed to further validate the results of the survey and actively monitor virus diversity and titer through the queen rearing process under controlled conditions in the research apiary at the DREC, Stoneville, MS, from May to September 2023. A single healthy, queen-right colony headed by a Pol-line queen [93] served as the sole source of larvae for grafting. Three double-deep standard Langstroth queenless colonies of Pol-line honey bees were also established as nurse colonies. The nurse colonies were densely populated with young worker bees, enabling them to care for the grafted larvae, produce and deposit sufficient royal jelly in each queen cell. Grafting was repeated three times over 5 weeks, and in each grafting event, we collected a pool of 20 eggs and newly hatched larvae to evaluate the load and diversity of viruses in the grafting sources. 72 hours post grafting, developing queen larvae were collected, where larvae and the corresponding royal jelly were sampled separately and stored in the -80 °C freezer for subsequent molecular analysis. While collecting queen larvae, a pool of ∼150 worker bees was collected from each nurse colony and stored in the -80 °C freezer to evaluate *Varroa* infestation level and viral infection. Subsequently, purple eye queen pupae were collected, and the remaining capped queen cells were transferred to an emergence incubator (35 °C and 70% RH) to let the queens emerge. Newly emerged queens were collected individually in microcentrifuge tubes, and the rest of the queens were marked and introduced to 5-frame mating nucs to mate naturally. Each queen was maintained in its original nucs until the termination of the experiment. Following the onset of egg laying, a pool of 20 freshly laid eggs, and a pool of worker bees were collected from each nuc. This sampling was repeated twice at an interval of 6 weeks. Soon after last egg sampling, all the queens from the mating nucs were collected in individual microcentrifuge tubes and stored in -80 °C freezer for molecular analysis. In total, 288 samples representing 10 sample types: pools of eggs (n=8) and larvae (n=6) from the mother colony, queen larvae (n=30), royal jelly (n=30), queen pupae (n=44), virgin queens (n=59), and pool of worker bees (n=11) from nurse colonies, and pool of eggs (n=40), pool of worker bees (n=40), and mated queens (n=20) from mating nucs, were collected for this experiment (S2 Table).

### Sample preparation and molecular assays

All the pooled samples of worker bees collected in the survey were assessed for *Varroa* infestation. From each pool of worker samples, 50 bees were randomly separated for molecular assay, individually checked for *Varroa* mites on a -20 °C surface. Since almost all mites could be harboring several bee viruses [11,94], they were removed from bees before RNA extraction to avoid overestimating the viral incidence and loads in bees. After removing mites from worker bees, samples were immediately preserved at -80 °C until RNA extraction. For the remaining worker bees, mites were counted using the alcohol (70% ethanol) wash method with a *Varroa* EasyCheck kit.

To initiate RNA extraction, 3 metal beads (2.4 mm, PerkinElmer) were added to each tube, and samples were homogenized using a Bead Ruptor Elite (OMNI International Inc.), as described previously [49]. Total RNA was extracted from each sample using the MagMAX mirVanna Total RNA Isolation Kit in a KingFisher Magnetic extractor (KingFisher Duo Prime, Thermo Fisher Scientific) following the manufacturer’s protocol with minor adaptation. The concentration and purity of the extracted RNA were measured using a NanoDrop One Microvolume UV-Vis Spectrophotometer (Thermo Fisher Scientific, MA, USA). The total RNA concentration was then adjusted to 50 ng/µL with the elution buffer provided in the extraction kit. Individual RNA samples were extracted and transferred into 96-well plates, then stored at - 80 ℃ for further use. A two-step PCR assay was performed to quantify a panel of 7 common honey bee viruses, including DWV-A, DWV-B (previously known as *Varroa destructor* virus-1), BQCV, SBV, IAPV, chronic bee paralysis virus (CBPV), and Lake Sinai viruses (LSVs). To do so, a High-Capacity cDNA Reverse-Transcription Kit (Applied Biosystems, Foster City, CA, USA) was used to synthesize cDNA. The reaction mixture was prepared by adding 10 μL of RNA template (500 ng) to 10 μL of the supplied cDNA master mix. The reaction was incubated as per the manufacturer’s instructions: 10 min at 25 °C, 120 min at 37 °C, and 5 min at 85 °C.

The resulting cDNA solution was then diluted 10-fold in nuclease-free water to serve as a template in subsequent qPCR.

The qPCR was conducted in duplicate using 384-well plates on a QuantStudio6™ Pro (Applied Biosystems) apparatus. The reactions were performed using previously validated primers (S3 Table) and the PowerUp SYBR Green master mix (Thermo Fisher Scientific Baltics UAB, Vilnius, Lithuania), in a total volume of 12 µL with a final primer concentration of 0.4 µM. RPS-5 was used as a reference gene to serve as an internal control to confirm the integrity of sample processing and functionality of the assay. On each plate, a ten-fold serial dilution of synthesized PCR products served as a positive control, RNase-free water as a template for a NO Target Control (NTC), and a No Reverse Transcriptase (NRT) control as an additional negative control [95]. The thermal cycling conditions were initialized by hold stage for 2 min at 50 °C and 95 °C, followed by 40 cycles consisting of a denaturation stage at 95 °C for 15 s and annealing/extension at 60 °C for 1 min, and a melt curve stage (95 °C for 15 sec, 60 °C for 1 min, and dissociation at 95 °C for 15 s). Fluorescence measurements were taken at the end of each cycle, and samples were considered positive for a target if the quantification cycle (Cq) value was recorded as 32 or lower. For analytical specificity, melt curves were analyzed to detect the formation of any primer dimers, and non-specific amplicons were discarded [95]. Viral load in each sample was quantified using absolute quantification based on standard curves obtained by serial dilutions of a known amount of viral amplification, as described before [20].

### Data handling and statistical analysis

All statistical analysis and visualization were performed using R version 4.4.1 [96] in RStudio version 2024.12.0+467 [97]. Data handling and organization were performed using the packages: readxl [98] and tidyverse [99]. *Varroa* mite infestation analysis was performed using the package glmmTMB [100]. Virus prevalence was calculated using the tidyverse package [99]. Model fitting and pairwise comparison for viral load analysis were performed using the packages: brms [101], bayestestR [102], and emmeans [103]. Hierarchical clustering was performed using the package pvclust [104]. For correlation analysis and table formatting, corr [105] and gt [106] packages were used. Plots were visualized using tidyverse [99], ggsignif [107], and ggpubr [108] packages.

The viral data obtained in both experiments (controlled and survey experiments) were divided into two subsets before statistical analysis. The first subset represents viral load in different developmental stages of queens in both experiments. The second subset represents viral load at the colony level in the survey, whereas in the controlled experiment, it represents viral load in eggs and worker bees from mating nucs, along with nurse bees. Statistical analysis was performed on the original scale of viral loads, except for the statistics of drone viral loads and the correlation between viral loads and mite level were presented in transformed viral loads. For better visualization, viral loads were log-transformed ((log_10_ virus copies+1) ng/RNA) prior to plotting.

To understand the distribution and diversity of viruses in the survey, virus prevalence was computed as the proportion of positive cases within each sample type. The normality and presence of outliers in the data were assessed using the Shapiro-Wilk test. Since the data were not normally distributed and contained a high frequency of zeroes (i.e., samples where no virus was detected), a generalized linear mixed model was fitted, with a family hurdle lognormal distribution. The model was fitted with Hamiltonian Monte Carlo and used an identity link for non-zero viral loads and a logit link (logistic regression) to estimate the likelihood of zero viral loads. We verified model convergence by ensuring the R-hat statistics for all parameters were below 1.01. The survey experiment contains different variables; therefore, the model was fitted with sample types representing viral loads across various queen developmental stages and worker bees in different colony types as fixed effects. Meanwhile, the queen breeders and the interaction between year and queen breeders were considered as random effects. Similarly, for the controlled experiment, the sample types were considered as the fixed effect, and the sample ID was taken as a random effect to account for non-independence among repeated measures within individual samples. Pairwise contrast was performed to compare viral loads across different sample types, including queen development stages and colonies for each of the seven viruses. We calculated the median and 95% equal-tailed credible interval for each estimated mean and ratio.

To explore the clustering patterns among different viral loads and sample types, hierarchical clustering was performed with multiple linkage methods. The clustering is based on the distance metric method, which measures pairwise dissimilarities among viral loads. As the viral loads failed to satisfy the assumptions for normality, Spearman’s correlation analysis was performed to assess the rank-based association between viral loads as well as between viral load and mite levels in different sample types. For the analysis of the *Varroa* mite infestation, a zero-inflated Gamma mixed model was used with the percentage of mite infestation as a fixed effect and queen breeder and the interaction between breeder and year as random effects.

## Acknowledgments

We wish to thank all the queen producers in Mississippi and Louisiana, who collaborated with us over two years of sampling, and generously provided access to their operations and donating samples that made this study possible. We also would like to thank Vanessa Bonatti from the Center for Pollinator Health at the Delta Research and Extension Center (DREC) for her assistance with part of the molecular analysis. We thank all members of the Center for Pollinator Health for their collegiality throughout this project.

## Supporting information

**S1 Fig. Varroa mite infestation level across colony types.** Each dot in the violin plot represents the percentage of mite infestation in a pool of worker bees. The shape of the violin plot represents the distribution density of mite infestation in each colony type. The different letters from post-hoc pairwise comparison indicate significant differences in mite infestation among colonies at a 5% level of significance. Breeder colonies exhibited the lowest mite infestation, whereas the nurse colony exhibited the highest mite infestation.

**S2 Fig. Correlation analysis between viral load in mated queens and Varroa mite infestation in the corresponding worker bees from mating nucs.** The correlation analysis was performed on these paired samples collected during the survey (2024) for DWV-A, DWV-B, BQCV, LSV, and CBPV, which showed no significant correlation between viral load and mite infestation. Correlation was assessed using Spearman correlation, in which R represents the correlation coefficient, while the p-value indicates the significance level of the correlation.

**S3 Fig. Correlation analysis between viral load in eggs and Varroa mite infestation in the corresponding worker bees from breeder colonies.** The correlation analysis was performed between these paired samples collected during the survey (2024) for DWV-A, DWV-B, BQCV, LSV, and CBPV, which showed no significant correlation between viral load and mite infestation. Correlation was assessed using Spearman correlation, in which R represents the correlation coefficient, while the p-value indicates the significance level of the correlation.

**S1 Table. Summary of samples collected during the survey (2023 & 2024).** Samples comprising both developmental stages of queens and worker bees from different colony types: breeder/mother colonies, nurse colonies, bank colonies, and mating nucs were collected. Worker bees were collected from only breeder and nurse colonies in 2023, while in 2024, they were collected from all the colony types. Therefore, 10 types of samples were collected in 2023, while it was 12 in 2024. The breeder (mother) colony is the source colony whose queen provides young larvae for producing new queens through grafting. Nurse colonies are used to rear developing queens after grafting young larvae, which are maintained with enough young nurse bees to take care of developing queens. Bank colonies are specially prepared colonies (either queenless or queenright), which are used to store queens in bulk for various purposes until required. Mating nucs or mating nuclei are the small honey bee colonies (often containing 3-5 frames) designed to support natural mating of virgin queens.

**S2 Table. Summary of samples collected during the controlled experiment.** Ten types of samples comprising both developmental stages of queens and worker bees were collected. Workers were sampled only from nurse colonies and mating nucs. The breeder (mother) colony is the source colony whose queen provides young larvae for producing new queens through grafting. Nurse colonies are used to rear developing queens after grafting young larvae, which are maintained with enough young nurse bees to take care of developing queens. Mating nucs or mating nuclei are the small honey bee colonies (often containing 3-5 frames) designed to support natural mating of virgin queens.

**S3 Table. Primers used in this study for qPCR and calibration curve generation.**

**S4 Table. Summary statistics of viral loads (log_10_ + 1) in Drones.** The viral prevalence and loads were computed by analyzing 10 individual drones per queen breeder, which were collected from mating nucs in the survey. BQCV showed the highest prevalence (85.47%), followed by DWV-B (70.39%). The prevalence of other viruses was below 50%. Regarding viral loads, BQCV exhibited the highest average titer (log_10_ (BQCV copies+1)/ng RNA = 4), followed by DWV-B and LSVs. Other viruses showed less than 1 log_10_ (virus copies+1)/ng RNA.

## References

1. Winston ML. The biology of the honey bee. Harvard University Press; 1991.

2. Winston ML, Slessor KN. Honey bee primer pheromones and colony organization: gaps in our knowledge Apis mellifera / primer pheromones / queen pheromones / colony integration. 1998.

3. Hoover SER, Keeling CI, Winston ML, Slessor KN. The effect of queen pheromones on worker honey bee ovary development. Naturwissenschaften. 2003;90: 477–480. doi:10.1007/s00114-003-0462-z

4. Beggs KT, Glendining KA, Marechal NM, Vergoz V, Nakamura I, Slessor KN, et al. Queen pheromone modulates brain dopamine function in worker honey bees. 2007. Available: 10.1073/pnas.0608224104

5. Oldroyd BP, Rinderer TE, Harbo JR, Buco SM. Effects of Intracolonial Genetic Diversity on Honey Bee (Hymenoptera: Apidae) Colony Performance. Ann Entomol Soc Am. 1992;85: 335–343. doi:10.1093/aesa/85.3.335

6. Rangel J, Keller JJ, Tarpy DR. The effects of honey bee (*Apis mellifera* L.) queen reproductive potential on colony growth. Insectes Soc. 2013;60: 65–73. doi:10.1007/s00040-012-0267-1

7. Yu L, Shi X, He X, Zeng Z, Yan W, Wu X. High-Quality Queens Produce High-Quality Offspring Queens. Insects. 2022;13: 486. doi:10.3390/insects13050486

8. Gauthier L, Ravallec M, Tournaire M, Cousserans F, Bergoin M, Dainat B, et al. Viruses associated with ovarian degeneration in *Apis mellifera* L. Queens. PLoS One. 2011;6. doi:10.1371/journal.pone.0016217

9. Amiri E, Strand MK, Rueppell O, Tarpy DR. Queen quality and the impact of honey bee diseases on queen health: Potential for interactions between two major threats to colony health. Insects. MDPI AG; 2017. doi:10.3390/insects8020048

10. Chapman A, McAfee A, Tarpy DR, Fine J, Rempel Z, Peters K, et al. Common viral infections inhibit egg laying in honey bee queens and are linked to premature supersedure. Sci Rep. 2024;14. doi:10.1038/s41598-024-66286-5

11. Chen YP, Higgins JA, Feldlaufer MF. Quantitative real-time reverse transcription-PCR analysis of deformed wing virus infection in the honeybee (*Apis mellifera* L.). Appl Environ Microbiol. 2005;71: 436–441. doi:10.1128/AEM.71.1.436-441.2005

12. Domingues CEC, Šimenc L, Toplak I, de Graaf DC, De Smet L, Verbeke W, et al. Eggs sampling as an effective tool for identifying the incidence of viruses in honey bees involved in artificial queen rearing. Sci Rep. 2024;14. doi:10.1038/s41598-024-60135-1

13. Francis RM, Nielsen SL, Kryger P. Patterns of viral infection in honey bee queens. Journal of General Virology. 2013;94: 668–676. doi:10.1099/vir.0.047019-0

14. Orlova M, harwood G, Freitak D, Amdam G. Immune challenge reduces the production of queen-specific compounds and fertility signals in honey bee queens. 2023. doi:10.21203/rs.3.rs-3221736/v1

15. Amiri E, Strand MK, Tarpy DR, Rueppell O. Honey bee queens and virus infections. Viruses. MDPI AG; 2020. doi:10.3390/v12030322

16. Shen M, Cui L, Ostiguy N, Cox-Foster D. Intricate transmission routes and interactions between picorna-like viruses (Kashmir bee virus and sacbrood virus) with the honeybee host and the parasitic varroa mite. Journal of General Virology. 2005;86: 2281–2289. doi:10.1099/vir.0.80824-0

17. Chen YP, Pettis JS, Collins A, Feldlaufer MF. Prevalence and transmission of honeybee viruses. Appl Environ Microbiol. 2006;72: 606–611. doi:10.1128/AEM.72.1.606-611.2006

18. Žvokelj L, Bakonyi T, Korošec T, Gregorc A. Appearance of acute bee paralysis virus, black queen cell virus and deformed wing virus in Carnolian honey bee (*Apis mellifera* carnica) queen rearing. J Apic Res. 2020;59: 53–58. doi:10.1080/00218839.2019.1681115

19. Yue C, Schröder M, Gisder S, Genersch E. Vertical-transmission routes for deformed wing virus of honeybees (*Apis mellifera*). Journal of General Virology. 2007;88: 2329–2336. doi:10.1099/vir.0.83101-0

20. Amiri E, Kryger P, Meixner MD, Strand MK, Tarpy DR, Rueppell O. Quantitative patterns of vertical transmission of deformed wing virus in honey bees. PLoS One. 2018;13. doi:10.1371/journal.pone.0195283

21. Amiri E, Meixner M, Büchler R, Kryger P. Chronic bee paralysis virus in honeybee queens: Evaluating susceptibility and infection routes. Viruses. 2014;6: 1188–1201. doi:10.3390/v6031188

22. Harizanis PC. Infestation of queen cells by the mite Varroa jacobsoni. 1991.

23. Bailey L, Woods RD. Two More Small RNA Viruses from Honey Bees and Further Observations on Sacbrood and Acute Bee-Paralysis Viruses. 1977.

24. Colwell MJ, CRW, PSF. Viruses in unexpected places: new transmission routes of European honey bee (*Apis mellifera*) viruses. American Bee Research Conference, Baton Rouge, LA. 2017; 106–107.

25. Chen Y, Pettis JS, Feldlaufer MF. Detection of multiple viruses in queens of the honey bee *Apis mellifera* L. J Invertebr Pathol. 2005;90: 118–121. doi:10.1016/j.jip.2005.08.005

26. Gregorc A, Bakonyi T. Viral infections in queen bees (*Apis mellifera* carnica) from rearing apiaries. Acta Veterinaria Brno. 2012;81: 15–19. doi:10.2754/avb201281010015

27. Yañez O, Jaffé R, Jarosch A, Fries I, Robin FAM, Robert JP, et al. Deformed wing virus and drone mating flights in the honey bee (*Apis mellifera*): Implications for sexual transmission of a major honey bee virus. Apidologie. 2012;43: 17–30. doi:10.1007/s13592-011-0088-7

28. Amiri E, Meixner MD, Kryger P. Deformed wing virus can be transmitted during natural mating in honey bees and infect the queens. Sci Rep. 2016;6. doi:10.1038/srep33065

29. Amiri E, Seddon G, Smith WZ, Strand MK, Tarpy DR, Rueppell O. Israeli acute paralysis virus: Honey bee queen–worker interaction and potential virus transmission pathways. Insects. 2019;10. doi:10.3390/insects10010009

30. Reid M. Storage of Queen Honeybees. Bee World. 1975;56: 21–31. doi:10.1080/0005772x.1975.11097534

31. Johnstone M. Rearing queen bees. 2008. Available: www.aqis.gov.au

32. Steinhaus EA. The importance of environmental factors in the insect-microbe ecosystem. Bacteriol Rev. 1960;24: 365–373.

33. Tlak Gajger I, Abou-Shaara HF, Smodiš Škerl MI. Strategies to Mitigate the Adverse Impacts of Viral Infections on Honey Bee (*Apis mellifera* L.) Colonies. Insects. 2025;16: 509. doi:10.3390/insects16050509

34. Tantillo G, Bottaro M, Di Pinto A, Martella V, Di Pinto P, Terio V. Virus infections of honeybees *Apis mellifera*. Italian Journal of Food Safety. Page Press Publications; 2015. pp. 157–168. doi:10.4081/ijfs.2015.5364

35. Lang S, Simone-Finstrom M, Healy K. Effects of honey bee queen exposure to deformed wing virus-A on queen and juvenile infection and colony strength metrics. J Apic Res. 2023;63: 233–244. doi:10.1080/00218839.2023.2284034

36. Chen YP, Pettis JS, Corona M, Chen WP, Li CJ, Spivak M, et al. Israeli Acute Paralysis Virus: Epidemiology, Pathogenesis and Implications for Honey Bee Health. PLoS Pathog. 2014;10. doi:10.1371/journal.ppat.1004261

37. Ravoet J, De Smet L, Wenseleers T, de Graaf DC. Vertical transmission of honey bee viruses in a Belgian queen breeding program. BMC Vet Res. 2015;11. doi:10.1186/s12917-015-0386-9

38. Blanchard P, Ribière M, Celle O, Lallemand P, Schurr F, Olivier V, et al. Evaluation of a real-time two-step RT-PCR assay for quantitation of Chronic bee paralysis virus (CBPV) genome in experimentally-infected bee tissues and in life stages of a symptomatic colony. J Virol Methods. 2007;141: 7–13. doi:10.1016/j.jviromet.2006.11.021

39. Ribière M, Ball B, Aubert M. Natural history and geographical distribution of honey bee viruses. Virology and the honey bee. 2008; 15–84.

40. Ribière M, Olivier V, Blanchard P. Chronic bee paralysis: A disease and a virus like no other? J Invertebr Pathol. 2010;103. doi:10.1016/j.jip.2009.06.013

41. Spivak M, Danka RG. Perspectives on hygienic behavior in *Apis mellifera* and other social insects. Apidologie. 2021;52: 1–16. doi:10.1007/s13592-020-00784-z

42. Bull JJ, Molineux IJ, Rice WR. Selection of benevolence in a host–parasite system. Evolution (N Y). 1991;45: 875–882. doi:10.1111/j.1558-5646.1991.tb04356.x

43. Bull JJ. Virulence. Evolution (N Y). 1994;48: 1423–1437. doi:10.1111/j.1558-5646.1994.tb02185.x

44. Cross ST, Maertens BL, Dunham TJ, Rodgers CP, Brehm AL, Miller MR, et al. Partitiviruses Infecting Drosophila melanogaster and Aedes aegypti Exhibit Efficient Biparental Vertical Transmission. J Virol. 2020;94. doi:10.1128/JVI.01070-20

45. Gauthier L, Tentcheva D, Tournaire M, Dainat B, Cousserans F, Colin ME, et al. Viral load estimation in asymptomatic honey bee colonies using the quantitative RT-PCR technique. Apidologie. 2007;38: 426–435. doi:10.1051/apido:2007026

46. Chen YP, Siede R. Honey Bee Viruses. 2007. pp. 33–80. doi:10.1016/S0065-3527(07)70002-7

47. Anderson DL. Pathogens and queen bees. Australasian Beekeeper. Australasian Beekeeper. 1993;94: 292–296.

48. Hashimoto Y, Valles SM. Infection characteristics of Solenopsis invicta virus 2 in the red imported fire ant, *Solenopsis invicta*. J Invertebr Pathol. 2008;99: 136–140. doi:10.1016/j.jip.2008.06.006

49. Valizadeh B, Hardy J, Chen J, Amiri E. Dynamics of Israeli acute paralysis virus in the red imported fire ant (*Solenopsis invicta* Buren) colonies. J Invertebr Pathol. 2025;211: 108310. doi:10.1016/j.jip.2025.108310

50. Mikhailov VS, Zemskov EA, Abramova EB. Protein synthesis in pupae of the silkworm *Bombyx mori* after infection with nuclear polyhedrosis virus: resistance to viral infection acquired during pupal period. Journal of General Virology. 1992;73: 3195–3202. doi:10.1099/0022-1317-73-12-3195

51. da Cruz-Landim C, Cavalcante VM. Ultrastructural and Cytochemical Aspects of Metamorphosis in the Midgut of *Apis mellifera* L. (Hymenoptera: Apidae: Apinae). Zoolog Sci. 2003;20: 1099–1107. doi:10.2108/zsj.20.1099

52. Remnant EJ, Mather N, Gillard TL, Yagound B, Beekman M. Direct transmission by injection affects competition among RNA viruses in honeybees. Proceedings of the Royal Society B: Biological Sciences. 2019;286: 20182452. doi:10.1098/rspb.2018.2452

53. Tehel A, Vu Q, Bigot D, Gogol-Döring A, Koch P, Jenkins C, et al. The Two Prevalent Genotypes of an Emerging Infectious Disease, Deformed Wing Virus, Cause Equally Low Pupal Mortality and Equally High Wing Deformities in Host Honey Bees. Viruses. 2019;11: 114. doi:10.3390/v11020114

54. Bailey L, Ball B V., Perry JN. Honeybee Paralysis: Its Natural Spread and its Diminished Incidence in England and Wales. J Apic Res. 1983;22: 191–195. doi:10.1080/00218839.1983.11100586

55. Chen Y, Evans J, Feldlaufer M. Horizontal and vertical transmission of viruses in the honey bee, *Apis mellifera*. J Invertebr Pathol. 2006;92: 152–159. doi:10.1016/j.jip.2006.03.010

56. Yue C, Schröder M, Bienefeld K, Genersch E. Detection of viral sequences in semen of honeybees (Apis mellifera): Evidence for vertical transmission of viruses through drones. J Invertebr Pathol. 2006;92: 105–108. doi:10.1016/j.jip.2006.03.001

57. Fievet J, Tentcheva D, Gauthier L, de Miranda J, Cousserans F, Colin ME, et al. Localization of deformed wing virus infection in queen and drone *Apis mellifera* L. Virol J. 2006;3: 16. doi:10.1186/1743-422X-3-16

58. Phokasem P, Liuhao W, Panjad P, Yujie T, Li J, Chantawannakul P. Differential Viral Distribution Patterns in Reproductive Tissues of *Apis mellifera* and *Apis cerana* Drones. Front Vet Sci. 2021;8. doi:10.3389/fvets.2021.608700

59. Gwynn DM, Callaghan A, Gorham J, Walters KFA, Fellowes MDE. Resistance is costly: trade-offs between immunity, fecundity and survival in the pea aphid. Proceedings of the Royal Society B: Biological Sciences. 2005;272: 1803–1808. doi:10.1098/rspb.2005.3089

60. Castella G, Christe P, Chapuisat M. Mating triggers dynamic immune regulations in wood ant queens. J Evol Biol. 2009;22: 564–570. doi:10.1111/j.1420-9101.2008.01664.x

61. Bascuñán-García AP, Lara C, Córdoba-Aguilar A. Immune investment impairs growth, female reproduction and survival in the house cricket, *Acheta domesticus*. J Insect Physiol. 2010;56: 204–211. doi:10.1016/j.jinsphys.2009.10.005

62. McNamara KB, Wedell N, Simmons LW. Experimental evolution reveals trade-offs between mating and immunity. Biol Lett. 2013;9. doi:10.1098/rsbl.2013.0262

63. Schwenke RA, Lazzaro BP, Wolfner MF. Reproduction–Immunity Trade-Offs in Insects. Annu Rev Entomol. 2016;61: 239–256. doi:10.1146/annurev-ento-010715-023924

64. Jasper WC, Brutscher LM, Grozinger CM, Niño EL. Injection of seminal fluid into the hemocoel of honey bee queens (*Apis mellifera*) can stimulate post-mating changes. Sci Rep. 2020;10: 11990. doi:10.1038/s41598-020-68437-w

65. McAfee A, Chapman A, Pettis JS, Foster LJ, Tarpy DR. Trade-offs between sperm viability and immune protein expression in honey bee queens (*Apis mellifera*). Commun Biol. 2021;4: 48. doi:10.1038/s42003-020-01586-w

66. Metz BN, Molina-Marciales T, Strand MK, Rueppell O, Tarpy DR, Amiri E. Physiological trade-offs in male social insects: Interactions among infection, immunity, fertility, size, and age in honey bee drones. J Insect Physiol. 2024;159: 104720. doi:10.1016/j.jinsphys.2024.104720

67. Baer B, Armitage SAO, Boomsma JJ. Sperm storage induces an immunity cost in ants. Nature. 2006;441: 872–875. doi:10.1038/nature04698

68. Barribeau SM, Schmid-Hempel P. Sexual healing: mating induces a protective immune response in bumblebees. J Evol Biol. 2017;30: 202–209. doi:10.1111/jeb.12964

69. Kokko H, Klug H, Jennions MD, Shuker DM, Simmons LW. Mating systems. The evolution of insect mating systems. 2014.

70. Oku K, Price TAR, Wedell N. Does mating negatively affect female immune defences in insects? Animal Biology. 2019;69: 117–136. doi:10.1163/15707563-20191082

71. Rolff J, Siva-Jothy MT. Copulation corrupts immunity: A mechanism for a cost of mating in insects. Proceedings of the National Academy of Sciences. 2002;99: 9916–9918. doi:10.1073/pnas.152271999

72. Schwenke RA, Lazzaro BP. Juvenile Hormone Suppresses Resistance to Infection in Mated Female *Drosophila melanogaster*. Current Biology. 2017;27: 596–601. doi:10.1016/j.cub.2017.01.004

73. Apanius V. Stress and Immune Defense. 1998. pp. 133–153. doi:10.1016/S0065-3454(08)60363-0

74. Steinhaus EA. Crowding as a Possible Stress Factor in Insect Disease. Ecology. 1958;39: 503–514. doi:10.2307/1931761

75. Morgan KN, Tromborg CT. Sources of stress in captivity. Appl Anim Behav Sci. 2007;102: 262–302. doi:10.1016/j.applanim.2006.05.032

76. Scharf I, Stoldt M, Libbrecht R, Höpfner AL, Jongepier E, Kever M, et al. Social isolation causes downregulation of immune and stress response genes and behavioural changes in a social insect. Mol Ecol. 2021;30: 2378–2389. doi:10.1111/mec.15902

77. Bernasconi G, Bigler L, Hesse M, Ratnieks FLW. Characterization of queen-specific components of the fluid released by fighting honey bee queens. Chemoecology. 1999;9: 161–167. doi:10.1007/s000490050049

78. Salas-Benito JS, De Nova-Ocampo M. Viral Interference and Persistence in Mosquito-Borne Flaviviruses. J Immunol Res. 2015;2015: 1–14. doi:10.1155/2015/873404

79. Goic B, Vodovar N, Mondotte JA, Monot C, Frangeul L, Blanc H, et al. RNA-mediated interference and reverse transcription control the persistence of RNA viruses in the insect model Drosophila. Nat Immunol. 2013;14: 396–403. doi:10.1038/ni.2542

80. Traynor KS, Pettis JS, Tarpy DR, Mullin CA, Frazier JL, Frazier M, et al. In-hive Pesticide Exposome: Assessing risks to migratory honey bees from in-hive pesticide contamination in the Eastern United States. Sci Rep. 2016;6: 33207. doi:10.1038/srep33207

81. D’Alvise P, Seeburger V, Gihring K, Kieboom M, Hasselmann M. Seasonal dynamics and co-occurrence patterns of honey bee pathogens revealed by high-throughput RT-qPCR analysis. Ecol Evol. 2019;9: 10241–10252. doi:10.1002/ece3.5544

82. Faurot-Daniels C, Glenny W, Daughenbaugh KF, McMenamin AJ, Burkle LA, Flenniken ML. Longitudinal monitoring of honey bee colonies reveals dynamic nature of virus abundance and indicates a negative impact of Lake Sinai virus 2 on colony health. PLoS One. 2020;15: e0237544. doi:10.1371/journal.pone.0237544

83. Spivak M. Honey bee hygienic behavior and defense against *Varroa jacobsoni*. Apidologie. 1996;27: 245–260. doi:10.1051/apido:19960407

84. Erez T, Bonda E, Kahanov P, Rueppell O, Wagoner K, Chejanovsky N, et al. Multiple benefits of breeding honey bees for hygienic behavior. J Invertebr Pathol. 2022;193: 107788. doi:10.1016/j.jip.2022.107788

85. Posada-Florez F, Childers AK, Heerman MC, Egekwu NI, Cook SC, Chen Y, et al. Deformed wing virus type A, a major honey bee pathogen, is vectored by the mite *Varroa destructor* in a non-propagative manner. Sci Rep. 2019;9: 12445. doi:10.1038/s41598-019-47447-3

86. Damayo JE, McKee RC, Buchmann G, Norton AM, Ashe A, Remnant EJ. Virus replication in the honey bee parasite, *Varroa destructor*. J Virol. 2023;97. doi:10.1128/jvi.01149-23

87. Ogihara MH, Behri M, Yoshiyama M. Detection of bee viruses from *Apis mellifera* (Hymenoptera: Apidae) and Varroa destructor (Acari: Varroidae) in Japan. Appl Entomol Zool. 2024;59: 293–303. doi:10.1007/s13355-024-00879-4

88. Kevill JL, Lee K, Goblirsch M, McDermott E, Tarpy DR, Spivak M, et al. The pathogen profile of a honey bee queen does not reflect that of her workers. Insects. 2020;11: 1–14. doi:10.3390/insects11060382

89. Grindrod I, Kevill JL, Villalobos EM, Schroeder DC, Martin SJ. Ten Years of Deformed Wing Virus (DWV) in Hawaiian Honey Bees (*Apis mellifera*), the Dominant DWV-A Variant Is Potentially Being Replaced by Variants with a DWV-B Coding Sequence. Viruses. 2021;13: 969. doi:10.3390/v13060969

90. Paxton RJ, Schäfer MO, Nazzi F, Zanni V, Annoscia D, Marroni F, et al. Epidemiology of a major honey bee pathogen, deformed wing virus: potential worldwide replacement of genotype A by genotype B. Int J Parasitol Parasites Wildl. 2022;18: 157–171. doi:10.1016/j.ijppaw.2022.04.013

91. Hesketh-Best PJ, Mckeown DA, Christmon K, Cook S, Fauvel AM, Steinhauer NA, et al. Dominance of recombinant DWV genomes with changing viral landscapes as revealed in national US honey bee and varroa mite survey. Commun Biol. 2024;7: 1623. doi:10.1038/s42003-024-07333-9

92. Sircoulomb F, Dubois E, Schurr F, Lucas P, Meixner M, Bertolotti A, et al. Genotype B of deformed wing virus and related recombinant viruses become dominant in European honey bee colonies. Sci Rep. 2025;15: 4804. doi:10.1038/s41598-025-86937-5

93. Danka RG, Harris JW, Dodds GE. Selection of VSH-derived “Pol-line” honey bees and evaluation of their Varroa-resistance characteristics. Apidologie. 2016;47: 483–490. doi:10.1007/s13592-015-0413-7

94. Tentcheva D, Gauthier L, Jouve S, Canabady-Rochelle L, Dainat B, Cousserans F, et al. Polymerase Chain Reaction detection of deformed wing virus (DWV) in *Apis mellifera* and *Varroa destructor*. Apidologie. 2004;35: 431–439. doi:10.1051/apido:2004021

95. Bustin SA, Benes V, Garson JA, Hellemans J, Huggett J, Kubista M, et al. The MIQE Guidelines: Minimum Information for Publication of Quantitative Real-Time PCR Experiments. Clin Chem. 2009;55: 611–622. doi:10.1373/clinchem.2008.112797

96. R Core Team. R: A language and environment for statistical computing. R Foundation for Statistical Computing, Vienna, Austria. 2024.

97. Posit team. RStudio: Integrated Development for RStudio, Inc., Boston, MA. 2024.

98. Wickham H, Bryan J. readxl: Read Excel Files. R package version 1.4.5. 2025. doi:https://readxl.tidyverse.org/

99. Wickham H, Averick M, Bryan J, Chang W, McGowan L, François R, et al. Welcome to the Tidyverse. J Open Source Softw. 2019;4: 1686. doi:10.21105/joss.01686

100. Brooks ME, Kristensen K, Benthem KJ, van, Magnusson A, Berg CW, Nielsen A, et al. glmmTMB Balances Speed and Flexibility Among Packages for Zero-inflated Generalized Linear Mixed Modeling. R J. 2017;9: 378. doi:10.32614/RJ-2017-066

101. Bürkner P-C. Advanced Bayesian Multilevel Modeling with the R Package brms. 2017. Available: http://arxiv.org/abs/1705.11123

102. Makowski D, Ben-Shachar M, Lüdecke D. bayestestR: Describing Effects and their Uncertainty, Existence and Significance within the Bayesian Framework. J Open Source Softw. 2019;4: 1541. doi:10.21105/joss.01541

103. Lenth R. emmeans: Estimated Marginal Means, aka Least-Squares Means. R package version 1.11.1. 2025.

104. Suzuki R, Shimodaira H. Pvclust: An R package for assessing the uncertainty in hierarchical clustering. Bioinformatics. 2006;22: 1540–1542. doi:10.1093/bioinformatics/btl117

105. Kuhn M, Jackson S, Cimentada J. corrr: Correlations in R. 2022.

106. Iannone R, Cheng J, Schloerke B, Hughes E, Lauer A, Seo J, et al. gt: Easily Create Presentation-Ready Display Tables. R package version 1.0.0.9000. 2025.

107. Ahlmann-Eltze C, Patil I. ggsignif: R Package for Displaying Significance Brackets for “ggplot2.” 2021.

108. Kassambara A. ggpubr: “ggplot2” Based Publication Ready Plots. R package version 0.6.0. 2023.

